# Ibex: Variational autoencoder for single-cell BCR sequencing

**DOI:** 10.1101/2022.11.09.515787

**Authors:** Nicholas Borcherding, Bo Sun, David DeNardo, Jonathan R. Brestoff

## Abstract

**Summary:** B cells are critical for adaptive immunity and are governed by the recognition of an antigen by the B cell receptor (BCR), a process that drives a coordinated series of signaling events and modulation of various transcriptional programs. Single-cell RNA sequencing with paired BCR profiling could offer insights into numerous physiological and pathological processes. However, unlike the plethora of single-cell RNA analysis pipelines, computational tools that utilize single-cell BCR sequences for further analyses are not yet well developed. Here we report Ibex, which vectorizes the amino acid sequence of the complementarity-determining region 3 (cdr3) of the immunoglobulin heavy and light chains, allowing for unbiased dimensional reduction of B cells using their BCR repertoire. Ibex is implemented as an R package with integration into both the Seurat and Single-Cell Experiment framework, enabling the incorporation of this new analytic tool into many single-cell sequencing analytic workflows and multimodal experiments.

**Availability and Implementation:** Ibex is available as an R package at https://github.com/ncborcherding/Ibex. Reproducible code and data for the figure appearing in the manuscript are available at https://github.com/ncborcherding/Ibex.manuscript. A companion TCR-based approach is available at https://github.com/ncborcherding/Trex.

## Introduction

Single-cell sequencing has become a mainstay in immunology. In addition to enabling the characterization of various immune populations and their transcriptional profiles, single-cell sequencing technologies allow for the pairing of transcriptomic and clonotypic data and can therefore provide insight into the complexities of the adaptive immune response. In the context of lymphocytes, the immune response is regulated by antigen recognition at the site of the adaptive immune receptor. In B cells, the BCR sequence is determined by the somatic recombination genes at heavy and light chain loci (Schatz and Ji, 2011) and in the germinal center via somatic hypermutation.(Odegard and Schatz, 2006) This process collectively produces groups of “clonal lineages” sequences that are generally used as categorical data in single-cell analysis.

Several tools have been put forth to integrate single-cell transcriptomic and T-cell receptor sequences, such as CoNGA (Schattgen *et al*., 2022), TESSA (Zhang *et al*., 2021), and mvTCR (Drost *et al*., 2022). The latter two tools leverage neural networks to generate a latent representation of the cdr3 sequence. The power of this approach has been shown in cancer detection (Beshnova *et al*., 2020) and in predicting responses to immunotherapy (Sidhom *et al*., 2022). Recent deep-learning approaches have shown promise in optimizing antibody selection (Mason *et al*., 2021). Other groups propose the potential of using natural language processing to better understand the linguistics of immunoglobulins (Ig) and improve in silico development(Vu *et al*., 2022; Ostrovsky-Berman *et al*., 2021). However, with the exception of the recent Benisse python package (Zhang *et al*., 2022), tools for integrating multimodal single-cell sequencing data with BCR sequences are lacking and nonexistent for the R environment. Using variational autoencoder models, we developed an intuitive tool to encode single-cell BCR light and heavy chain sequences. For each chain, the cdr3 sequence is directly converted to a format for one-hot autoencoding or first converted into an amino acid property matrix, such as Atchley factors (Atchley *et al*., 2005) or Kidera factors (Kidera *et al*., 1985) This latter conversion has been applied to TCR strategies, such in motif-based repertoire profiling via support vector machines.(Thomas *et al*., 2014). The respective chain- and approach-specific model is then applied to the resulting transformation to produce encoded numerical vectors. With the Ibex package, the encoded numerical vectors can be incorporated into single-cell objects and used directly for dimensional reduction or multimodal integration. This provides an uncomplicated and adaptable approach for the incorporation of BCR sequences to the analysis of single-cell B-cell data. In addition, Ibex is compatible with our previous scRepertoire R package (Borcherding *et al*., 2020), enabling users to easily attach adaptive immune receptor sequences to single-cell objects. To demonstrate the application of Ibex, we investigated single-cell peripheral blood B cells in COVID-19-associated multisystem inflammatory syndrome (Ramaswamy *et al*., 2021).

## Results

Ibex is an R package that translates the BCR light and heavy chain cdr3 sequences using variational autoencoders. Each available autoencoder was trained on 800,000 productive human cdr3 sequences of the respective Ig chain. To provide for more robust models, training and testing data cohorts were collected across a range of disease processes, including infection, autoimmunity, malignancy, and healthy controls (Supplemental Table 1). Sequence length for heavy (mean = 16.88, median = 17.00, sd = 3.71) and light chain (mean = 11.92, median = 12, sd =1.41). The transformer used a 128-64-30-64-128 neuron structure in the keras R package (v2.4.0). Differential network architecture was explored is available in Supplemental Figure 1. Each autoencoding training involved zero padding up to 70 residues to allow for variable cdr3 lengths. Hyperparameters for the models were based on minimal loss functions (Supplementary Figure 2), which led to the following: batch size of 64, learning rate of 0.001, and latent dimensions of 30. The optimization utilized a stochastic gradient descent approach known as rectified Adam based on performance and speed. Both Adam and RMSprop optimization were evaluated and not utilized. In addition, each model was trained using 80:20 data split and had early stopping monitoring for minimal value loss and patience of 10 epochs. The output of these autoencoder models is a 30-dimensional latent space representing the amino acid cdr3 sequence (Figure 1A).

**Figure 1.**
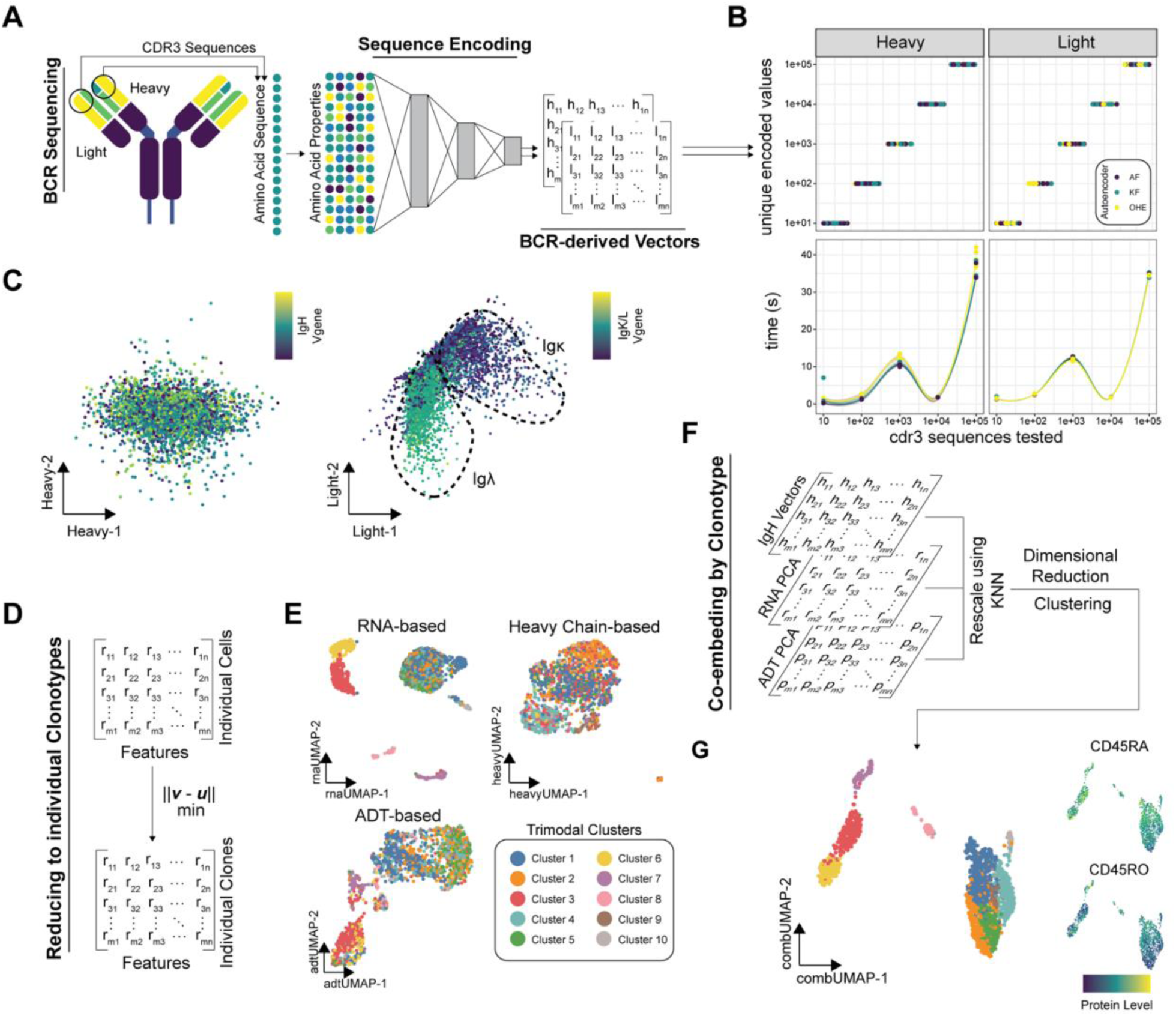
(**A**) Conceptual schematic for the autoencoding approach for the BCR sequences. (B) Benchmarking of the autoencoders in Ibex using novel cdr3 sequences for heavy and light chain in terms of unique latent dimensions returned (upper panels) and time for encoding (bottom panels). Colors are based on the encoding approach, Atchley factors (AF), Kidera factors (KF), and one-hot encoding (OHE). (**C**) Visualization of the latent space dimensions using the KF for the heavy (left) and light (right) chains. Colors indicate differing V genes associated with the respective chain. (**D**) Reduction of individual cells to individual clonotypes by selecting cells with the minimal Euclidean distance in principal component vectors across clonotypes. (**E**) Uniformed manifold approximation and projection (UMAP) using RNA, ADT-level protein quantification, and Ibex-derived heavy chain sequences. Clustering based on trimodal integrated values. (**F**) Approach for the multimodal integration of all three data types through k nearest neighbor rescaling. (**G**) Resulting UMAP from the normalized trimodal integrated values with protein quantification of naïve (CD45RA+) and antigen-experienced (CD45RO+) B cells.

To benchmark the autoencoding approach for both heavy and light chains, a bootstrapping approach using randomized novel cdr3 sequences that no model had seen (Figure 1B). For each model, 10, 100, 1000, 1e5, and 1e6 novel sequences were tested 10 times. Comparing the number of cdr3 sequences assessed to the unique latent dimensional output for the models demonstrated concordance for both the light and heavy sequences (Figure 1B, upper panel).

The encoding time between the approaches available in Ibex was relatively consistent, with encoding 1e6 sequences taking under 40 seconds of computation (Figure 1B, lower panel).

In order to demonstrate the application of Ibex, we selected a publicly-available data set of pediatric multisystem inflammatory syndrome patients (MIS) (Ramaswamy *et al*., 2021). This data set has the advantage of providing peripheral blood samples with single-cell RNA and protein quantifications with adaptive immune receptor sequencing. After combining the data from two MIS patients and two healthy controls from the cohort, B cells were isolated using canonical lineage markers and BCR sequences. After which, Ibex encoding for both the light and heavy chains of the B cells was performed by converting the amino acid sequence into Kidera factors. Latent dimensions and multidimensional scaling for all three models across the two chains are available in Supplementary Figure 3. The latent dimensions from the encoding can be used to directly explore sequence similarities, such as the clustering of IgK and Igλ (Figure 1C). The zero-padding of cdr3 up to 70 residues had minimal effect on latent dimensional embedding (Supplemental Figure 4), with potential skewing in longer cdr3 sequences. The skewing in the heavy chain may be due to the poor representation of longer sequences in the training data set (95% heavy sequences < 24.3 residues). Differential positions of light chain embedding were largely a factor of k/λ V gene usage. Ibex also enables the reduction of cell-level quantifications to clonotype-level quantifications using minimal Euclidean distance across principal component dimensions (Figure 1D), as previously described (Schattgen *et al*., 2022), or as an average across all cells defined as clones. The clonotype-level single-cell data can be used to generate UMAPs for the RNA, antibody-derived tag (ADT) protein level quantification, or the Ibex-based chain encoded values (Figure 1E). However, each of these assays can be rescaled across modalities to generate normalized values using packages such as mumosa R package (Lun, Aaron, 2022) (Figure 1F) or weighted-nearest neighbors in the Seurat R package (Hao *et al*., 2021). The resulting weighted or normalized values can then be used to create a unified representation of RNA, protein, and Ig heavy chain, such as with dimensional reduction or clustering (Figure 1G). We compared the outputs Ibex and Benisse (Zhang *et al*., 2022) for RNA and heavy chain cdr3 embedding (Supplemental Figure 5). The clustering from Benisse had higher granularity, which may result from the pipeline’s use of both nucleotide and amino acid sequences as input to the encoding model. We also found minimal overlap between the nearest neighbors called by the Ibex co-embedding and Benisse, suggesting these two approaches may have distinct strengths and could be complementary.

## Conclusion

Ibex is an R package designed to combine deep learning with immune repertoire profiling and incorporate them into common single-cell sequencing analytic workflows. The package offers customizable encoding of BCR sequences to produce latent dimensional representations of amino acid sequences. Unique to Ibex is the support of heavy and light chain vectorization across multiple deep learning models. Alone or in combination with other single-cell modalities, these latent vectors may assist in characterizing an immune response, understanding Ig maturation, or possible epitope identification. In future work, we will develop additional deep learning models or incorporate previously published approaches such as immuneML (Pavlović *et al*., 2021) or Immune2vec (Ostrovsky-Berman *et al*., 2021). Additionally, incorporating clonal lineage/mutation or the combination of sequence, expression, and mutational models into a deep learning model is particularly interesting. Distinct difference in the reconstruction of encoding methods between network architecture (Supplemental Figure 1) and encoding methods (Supplemental Figure 2) suggests there are additional areas of development for future models.

## Financial Support

This work was supported by the National Institutes of Health (NIH) Common Fund via the NIH Director’s Early Independence Award (DP5 OD028125) to J.R.B and Departmental Funding from the Department of Pathology and Immunology at Washington University.

## Conflicts of Interest

NB is a consultant for Santa Ana Bio, Inc and Omniscope, Inc. The other authors have no conflicts of interest relating to the work described in this manuscript.

## Notes

https://github.com/ncborcherding/Ibex.manuscript

https://github.com/ncborcherding/Ibex

### Competing Interest Statement

NB is a consultant for Santa Ana Bio, Inc and Omniscope, Inc. The results outlined in this publication are independent of these interests.

### Summary of Updates

Per reviewer comments, supplemental materials have been included to allow users to 1) evaluate use of models, 2) understand model development, and 3) compare to alternative pipelines.

