## Supplemental Materials for "Ibex: Variational autoencoder for single-cell BCR sequencing"

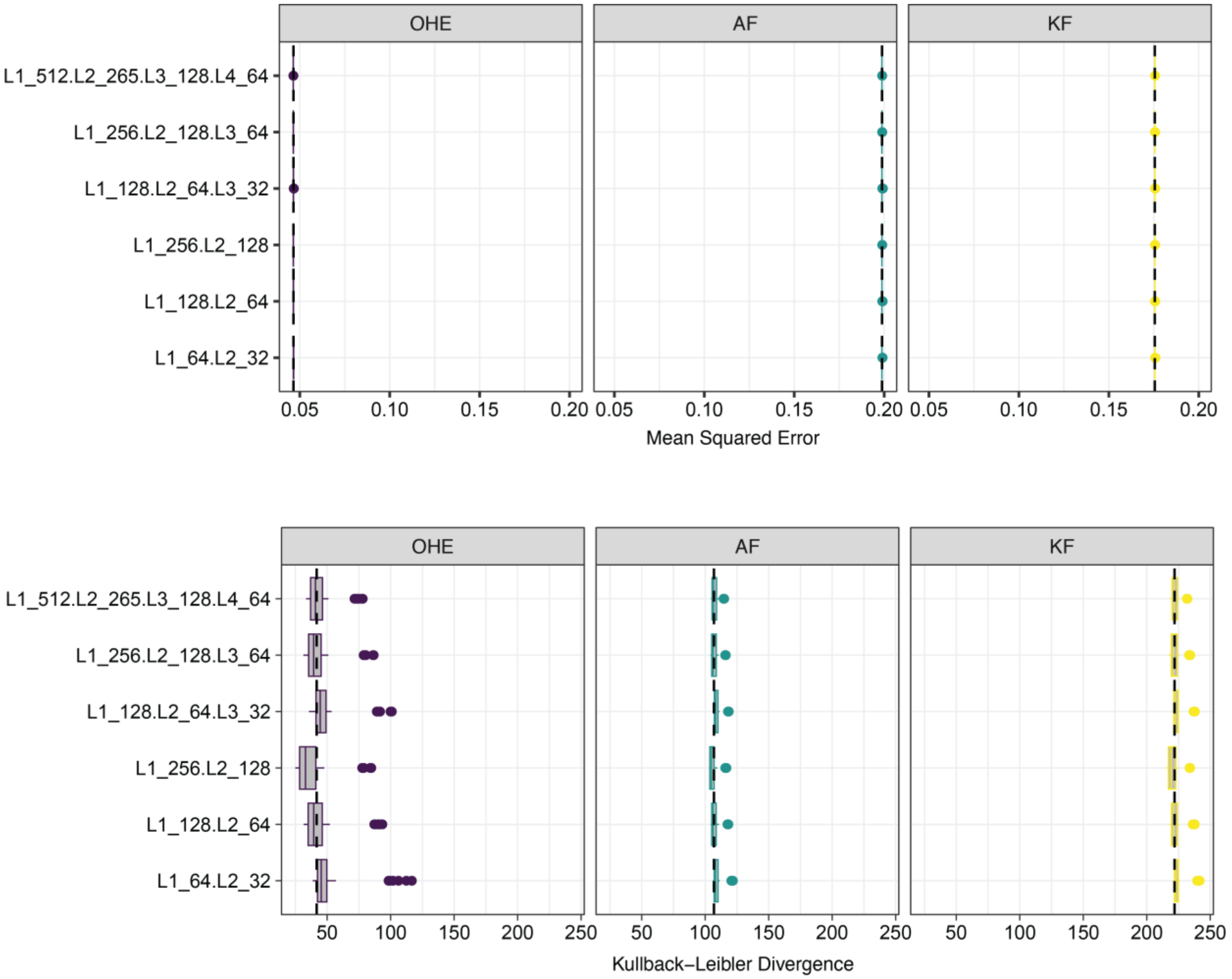


**Supplemental Figure 1:** Performance metrics for autoencoder models by modulating the transformer architecture. Mean square error values (upper panel) and Kullback-Leibler Divergence values (lower panel) for example heavy chain model training. Encoding models layers and number of neurons are indicated on the y-axis with L1_512 standing for layer 1, 512 neurons. A dotted line indicates the median value across models.

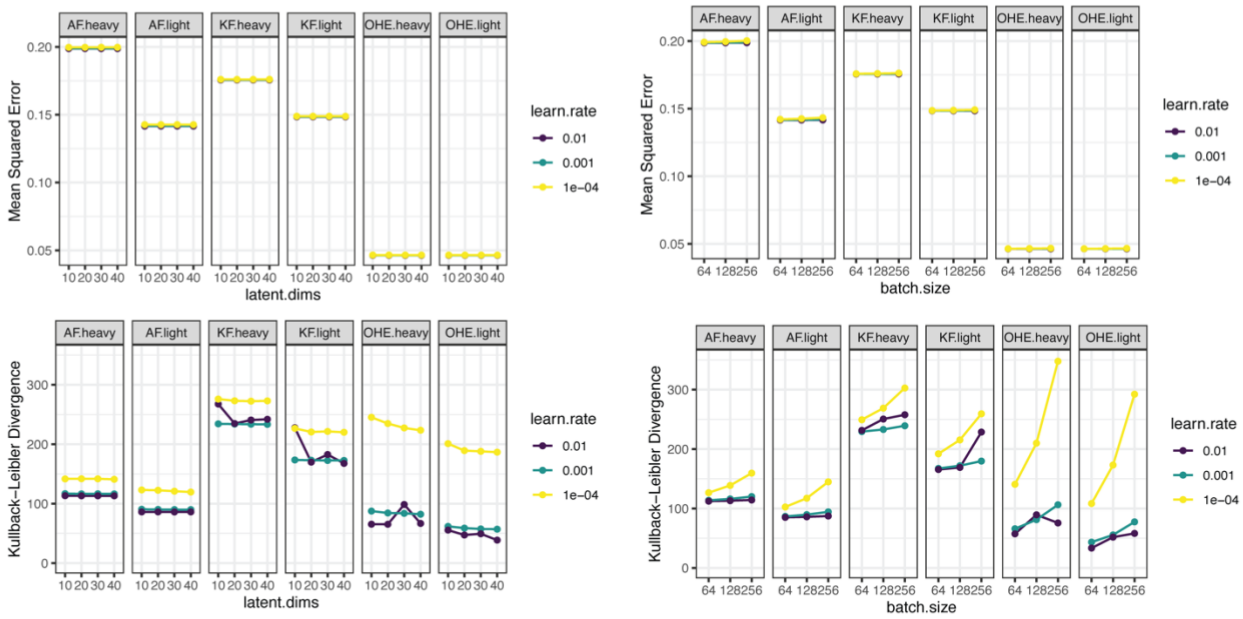


**Supplemental Figure 2:** Performance metrics for autoencoder models by approach and chain. For given hyperparameter, models were trained on 2e5 random sequences with 10 epochs for minimal Kullback-Leibler divergence value. Mean square error (upper panels) and Kullback-Leibler divergence values (lower panels) of models after training varying the latent dimensions (left panel) and batch size (right panel) with different learning rates.

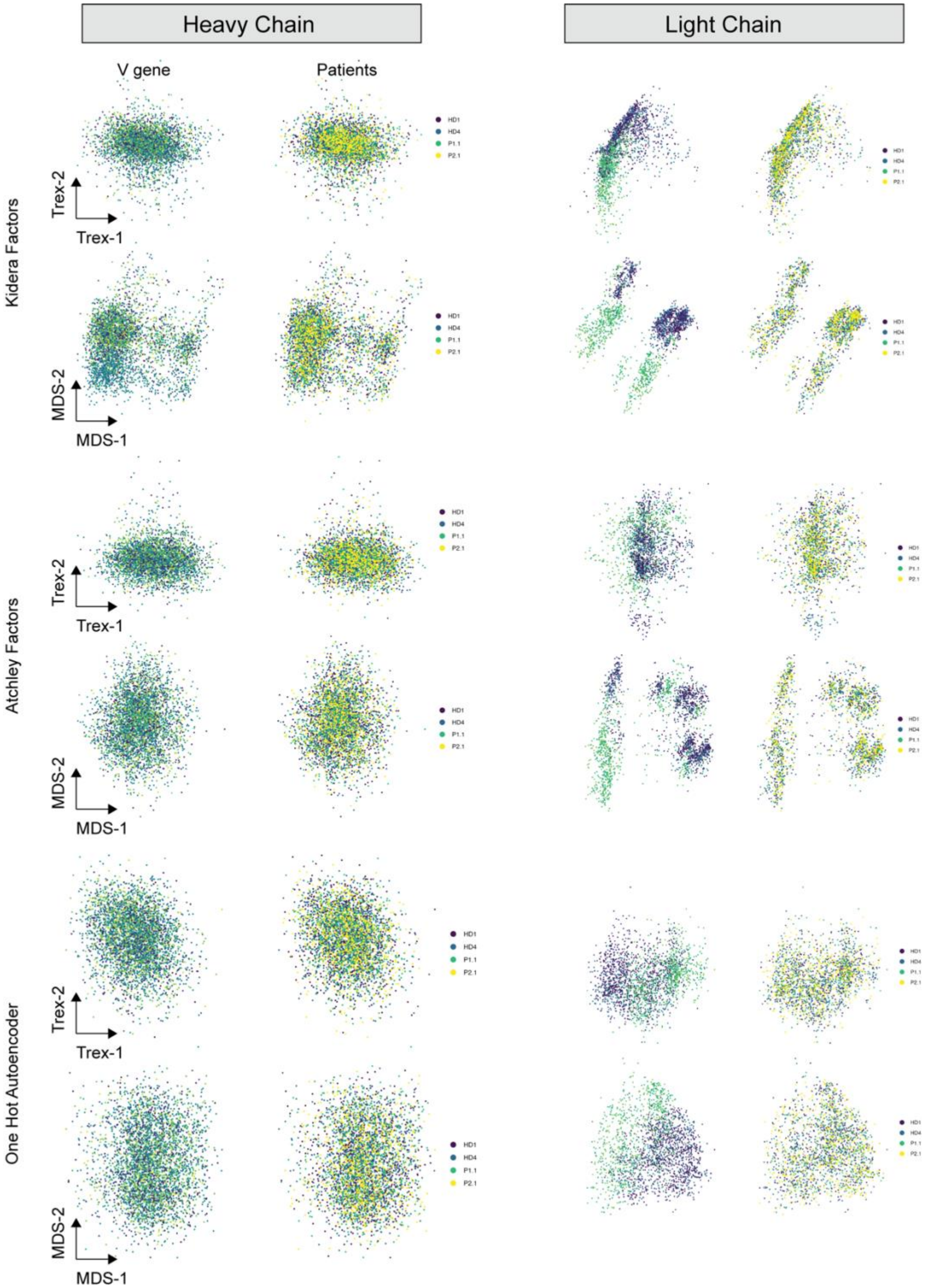


**Supplemental Figure 3:** Direct latent dimensional encoding and multidimensional scaling for models using KF, AF, and one-hot autoencoding for heavy (left column) and light chains (right column). Colors based on the respective v gene for the chain (left column) and the patient/donor (left column).

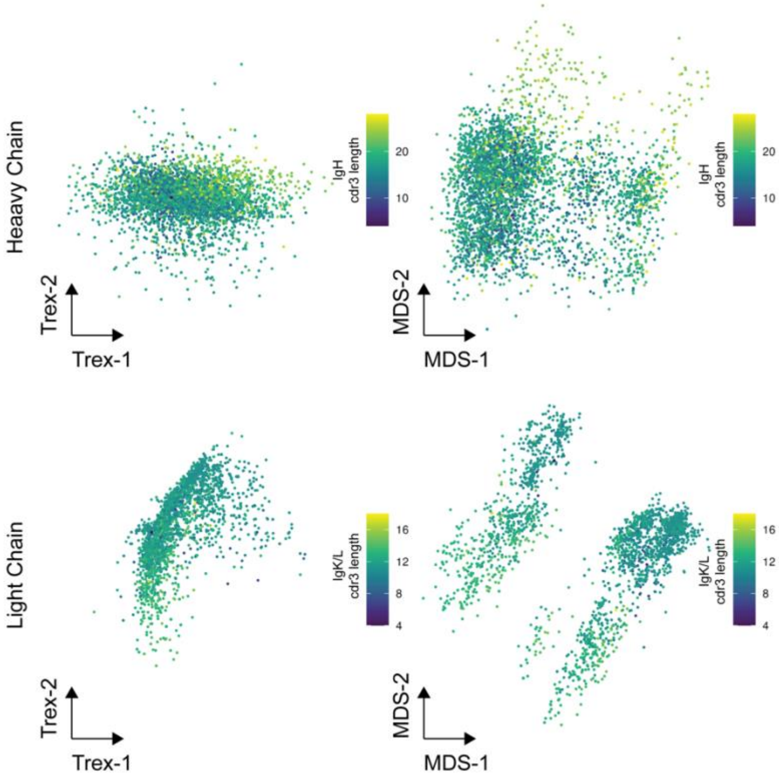


**Supplemental Figure 4:** Direct latent dimensional encoding and multidimensional scaling using the Kidera factors of the cdr3 amino acid sequences. Colors are based on the length of the respective sequence.

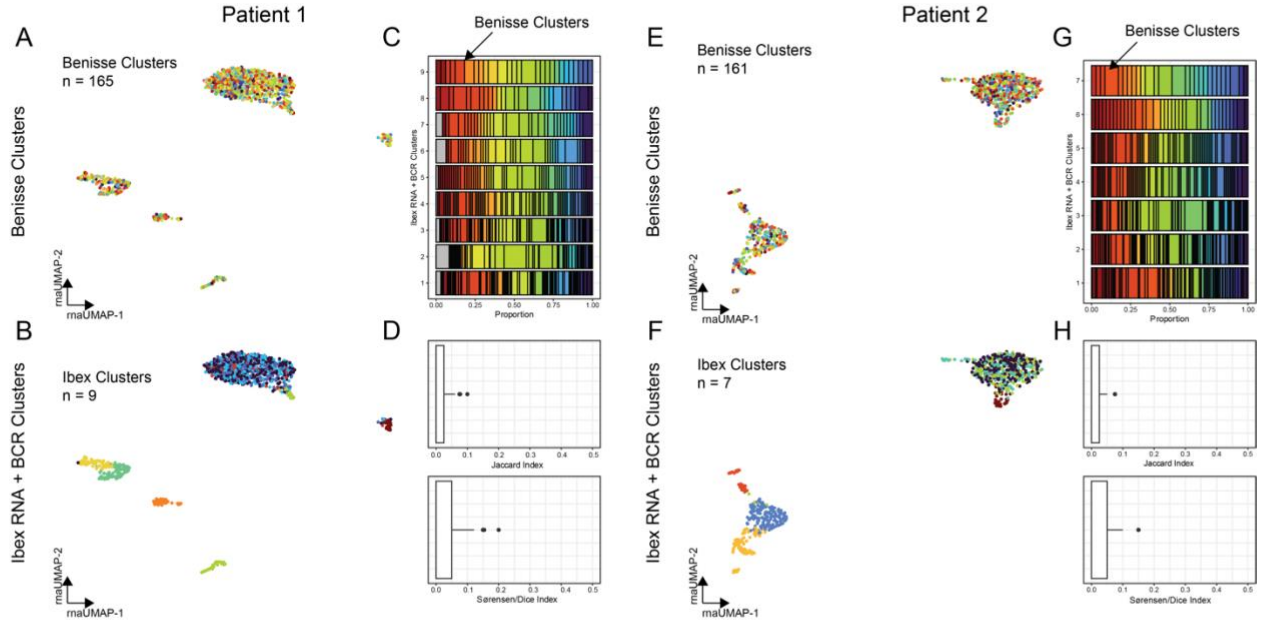


Supplemental Figure 5: Comparison of RNA and Heavy chain BCR embedding using Benisse and Ibex. The right set of panels are for MIS patient 1 and the left panel is for patient 2. A/E. An mRNA-based UMAP with Benisse cluster overlay. B/F. An mRNA-based UMAP with Ibex cluster overlay. C/G. The relative distribution of Ibex and Benisse. D/H. The Jaccard index and Sorensen-Dice index are based on the intersection of the 20 nearest neighbors from each method. Nearest neighbors for Benisse were isolated using the distance outputs from Benisse and the nearest-neighbor function in Seurat for Ibex co-embedding values.

Supplemental Table 1

| File_Name | Acession | Disease | Cite |
|---|---|---|---|
| 192561SKL_filtered_contig_annotations.csv | GSE169440 | Lyme Erythema Migrans | |
| 192561SKN_filtered_contig_annotations.csv | GSE169440 | Lyme Erythema Migrans | |
| 192563SKL_filtered_contig_annotations.csv | GSE169440 | Lyme Erythema Migrans | |
| 192564SKL_filtered_contig_annotations.csv | GSE169440 | Lyme Erythema Migrans | |
| 192564SKN_filtered_contig_annotations.csv | GSE169440 | Lyme Erythema Migrans | |
| 192565SKL_filtered_contig_annotations.csv | GSE169440 | Lyme Erythema Migrans | |
| 192565SKN_filtered_contig_annotations.csv | GSE169440 | Lyme Erythema Migrans | |
| 192566_filtered_contig_annotations.csv | GSE169440 | Lyme Erythema Migrans | |
| 192566SKL_filtered_contig_annotations.csv | GSE169440 | Lyme Erythema Migrans | |
| 192566SKN_filtered_contig_annotations.csv | GSE169440 | Lyme Erythema Migrans | |
| 192567SKL_filtered_contig_annotations.csv | GSE169440 | Lyme Erythema Migrans | |
| 192567SKN_filtered_contig_annotations.csv | GSE169440 | Lyme Erythema Migrans | [1] |
| 200142A_COV_144_filtered_contig_annotations.csv | GSE147507 | COVID-19 | |
| 200142A_COV_146_filtered_contig_annotations.csv | GSE147507 | COVID-19 | |
| 200142A_COV_147_filtered_contig_annotations.csv | GSE147507 | COVID-19 | |
| 200142A_COV_154_filtered_contig_annotations.csv | GSE147507 | COVID-19 | |
| 200142A_COV_155_filtered_contig_annotations.csv | GSE147507 | COVID-19 | |
| 200142A_COV_156_filtered_contig_annotations.csv | GSE147507 | COVID-19 | |
| 200142A_COV_157_filtered_contig_annotations.csv | GSE147507 | COVID-19 | |
| 200142A_COV_158_filtered_contig_annotations.csv | GSE147507 | COVID-19 | |
| 200142A_COV_159_filtered_contig_annotations.csv | GSE147507 | COVID-19 | |
| 200142A_COV_161_filtered_contig_annotations.csv | GSE147507 | COVID-19 | |
| 200142A_COV_163_filtered_contig_annotations.csv | GSE147507 | COVID-19 | |
| 200142A_HC_1_filtered_contig_annotations.csv | GSE147507 | COVID-19 | |
| 200142A_HC_05_filtered_contig_annotations.csv | GSE147507 | COVID-19 | |
| 200142A_HC_31_filtered_contig_annotations.csv | GSE147507 | COVID-19 | |
| 200142A_HC_36_filtered_contig_annotations.csv | GSE147507 | COVID-19 | |
| 200142B_COV_007_filtered_contig_annotations.csv | GSE147507 | COVID-19 | |
| 200142B_COV_009_filtered_contig_annotations.csv | GSE147507 | COVID-19 | |
| 200142B_COV_011_filtered_contig_annotations.csv | GSE147507 | COVID-19 | |
| 200142B_COV_012_filtered_contig_annotations.csv | GSE147507 | COVID-19 | |
| 200142B_COV_013_filtered_contig_annotations.csv | GSE147507 | COVID-19 | |
| 200142B_COV_016_filtered_contig_annotations.csv | GSE147507 | COVID-19 | |
| 200142B_COV_021_filtered_contig_annotations.csv | GSE147507 | COVID-19 | |
| 200142B_COV_029_filtered_contig_annotations.csv | GSE147507 | COVID-19 | |
| 200142B_COV_037_filtered_contig_annotations.csv | GSE147507 | COVID-19 | |
| 200142B_COV_045_filtered_contig_annotations.csv | GSE147507 | COVID-19 | |
| 200142B_COV_047_filtered_contig_annotations.csv | GSE147507 | COVID-19 | [2] |

| | | | |
|---|---|---|---|
| 200142B_COV_48_filtered_contig_annotations.csv | GSE147507 | COVID-19 | |
| 200142B_COV_055_filtered_contig_annotations.csv | GSE147507 | COVID-19 | |
| 200142B_COV_057_filtered_contig_annotations.csv | GSE147507 | COVID-19 | |
| 200142B_COV_072_filtered_contig_annotations.csv | GSE147507 | COVID-19 | |
| 200142B_COV_074_filtered_contig_annotations.csv | GSE147507 | COVID-19 | |
| 200142B_COV_76_filtered_contig_annotations.csv | GSE147507 | COVID-19 | |
| 200142B_COV_077_filtered_contig_annotations.csv | GSE147507 | COVID-19 | |
| 200142B_COV_078_filtered_contig_annotations.csv | GSE147507 | COVID-19 | |
| 200142B_COV_086_filtered_contig_annotations.csv | GSE147507 | COVID-19 | |
| 200142B_COV_87_filtered_contig_annotations.csv | GSE147507 | COVID-19 | |
| 200142B_COV_089_filtered_contig_annotations.csv | GSE147507 | COVID-19 | |
| 200142B_COV_91_filtered_contig_annotations.csv | GSE147507 | COVID-19 | |
| 200142B_COV_098_filtered_contig_annotations.csv | GSE147507 | COVID-19 | |
| 200142B_COV_101_filtered_contig_annotations.csv | GSE147507 | COVID-19 | |
| 200142B_COV_102_filtered_contig_annotations.csv | GSE147507 | COVID-19 | |
| 200142B_COV_119_filtered_contig_annotations.csv | GSE147507 | COVID-19 | |
| 200142B_COV_126_filtered_contig_annotations.csv | GSE147507 | COVID-19 | |
| 200142B_HC_07_filtered_contig_annotations.csv | GSE147507 | COVID-19 | |
| 200142B_HC_20_filtered_contig_annotations.csv | GSE147507 | COVID-19 | |
| 200142B_HC_21_filtered_contig_annotations.csv | GSE147507 | COVID-19 | |
| 200142B_HC_23_filtered_contig_annotations.csv | GSE147507 | COVID-19 | |
| 200142B_HC_39_filtered_contig_annotations.csv | GSE147507 | COVID-19 | |
| 200142B_HC_58_filtered_contig_annotations.csv | GSE147507 | COVID-19 | |
| 200142B_HC_60_filtered_contig_annotations.csv | GSE147507 | COVID-19 | |
| 200142C_COV_164_filtered_contig_annotations.csv | GSE147507 | COVID-19 | |
| 200142C_COV_165_filtered_contig_annotations.csv | GSE147507 | COVID-19 | |
| 200142C_COV_166_filtered_contig_annotations.csv | GSE147507 | COVID-19 | |
| GSE159929_filtered_contig_annotations_BCR.csv.gz | GSE159929 | Healthy | 3 |
| GSM4039806_165_filtered_contig_annotations.csv.gz | GSE136035 | SLE | |
| GSM4039808_178_filtered_contig_annotations.csv.gz | GSE136035 | SLE | |
| GSM4039810_182_filtered_contig_annotations.csv.gz | GSE136035 | SLE | |
| GSM4039814_177_filtered_contig_annotations.csv.gz | GSE136035 | Healthy | |
| GSM4885424_HC2_BCR_filtered_contig_annotations.csv.gz | GSE136035 | Healthy | |
| GSM4885426_HC3_BCR_filtered_contig_annotations.csv.gz | GSE136035 | Healthy | 4 |
| GSM5646528_tm1_BCR_filtered_contig_annotations.csv.gz | GSE186368 | Healthy | |
| GSM5646531_tm2_BCR_filtered_contig_annotations.csv.gz | GSE186368 | Healthy | |
| GSM5646534_tm3_BCR_filtered_contig_annotations.csv.gz | GSE186368 | Healthy | 5 |

| | | | |
|---|---|---|---|
| GSM5646537_tm4_BCR_filtered_contig_annotations.csv.gz | GSE186368 | Healthy | |
| GSM5646540_tm7_BCR_filtered_contig_annotations.csv.gz | GSE186368 | Healthy | |
| GSM5646543_tm8_BCR_filtered_contig_annotations.csv.gz | GSE186368 | Healthy | |
| GSM5694814_YUCARD_BCR.csv.gz | GSE189126 | Melanoma | |
| GSM5694815_YUENZO_BCR.csv.gz | GSE189126 | Melanoma | |
| GSM5694816_YUFUB_BCR.csv.gz | GSE189126 | Melanoma | |
| GSM5694817_YUFURL_BCR.csv.gz | GSE189126 | Melanoma | |
| GSM5694818_YUGRUS_BCR.csv.gz | GSE189126 | Melanoma | |
| GSM5694819_YUHERN_BCR.csv.gz | GSE189126 | Melanoma | |
| GSM5694820_YUKEND_BCR.csv.gz | GSE189126 | Melanoma | |
| GSM5694821_YUPIXEL_BCR.csv.gz | GSE189126 | Melanoma | |
| GSM5694822_YUROD_BCR.csv.gz | GSE189126 | Melanoma | |
| GSM5694823_YUTAUR_BCR.csv.gz | GSE189126 | Melanoma | |
| GSM5694824_YUTHEA_BCR.csv.gz | GSE189126 | Melanoma | |
| GSM5694825_YUTORY_BCR.csv.gz | GSE189126 | Melanoma | |
| GSM5694826_YUVARDO_BCR.csv.gz | GSE189126 | Melanoma | 6 |
| GSM5831588_PD1_filtered_contig_annotations.csv.gz | GSE194245 | Parkinsons Disease | |
| GSM5831589_PD2_filtered_contig_annotations.csv.gz | GSE194245 | Parkinsons Disease | |
| GSM5831590_PD3_filtered_contig_annotations.csv.gz | GSE194245 | Parkinsons Disease | |
| GSM5831591_PD4_filtered_contig_annotations.csv.gz | GSE194245 | Parkinsons Disease | |
| GSM5831592_PD5_filtered_contig_annotations.csv.gz | GSE194245 | Parkinsons Disease | |
| GSM5831593_PD6_filtered_contig_annotations.csv.gz | GSE194245 | Parkinsons Disease | |
| GSM5831594_PD7_filtered_contig_annotations.csv.gz | GSE194245 | Parkinsons Disease | |
| GSM5831595_PD8_filtered_contig_annotations.csv.gz | GSE194245 | Parkinsons Disease | |
| GSM5831596_HC1_filtered_contig_annotations.csv.gz | GSE194245 | Parkinsons Disease | |
| GSM5831597_HC2_filtered_contig_annotations.csv.gz | GSE194245 | Parkinsons Disease | |
| GSM5831598_HC3_filtered_contig_annotations.csv.gz | GSE194245 | Parkinsons Disease | |
| GSM5831599_HC4_filtered_contig_annotations.csv.gz | GSE194245 | Parkinsons Disease | |
| GSM5831600_HC5_filtered_contig_annotations.csv.gz | GSE194245 | Parkinsons Disease | |
| GSM5831601_HC6_filtered_contig_annotations.csv.gz | GSE194245 | Parkinsons Disease | 7 |
| GSM6341264_B1_all_contig_annotations.csv.gz | GSE208337 | COVID-19 | |
| GSM6341265_B2_all_contig_annotations.csv.gz | GSE208337 | COVID-19 | |
| GSM6341266_B3_all_contig_annotations.csv.gz | GSE208337 | COVID-19 | |
| GSM6341267_B4_all_contig_annotations.csv.gz | GSE208337 | COVID-19 | |
| GSM6341268_B5_all_contig_annotations.csv.gz | GSE208337 | COVID-19 | |
| GSM6341269_B6_all_contig_annotations.csv.gz | GSE208337 | COVID-19 | |
| GSM6341270_B7_all_contig_annotations.csv.gz | GSE208337 | COVID-19 | |
| GSM6341271_B8_all_contig_annotations.csv.gz | GSE208337 | COVID-19 | 8 |

| | | | |
|---|---|---|---|
| GSM6341272_B9_all_contig_annotations.csv.gz | GSE208337 | COVID-19 | |
| GSM6341273_B10_all_contig_annotations.csv.gz | GSE208337 | COVID-19 | |
| GSM6341274_B11_all_contig_annotations.csv.gz | GSE208337 | COVID-19 | |
| GSM6341275_B12_all_contig_annotations.csv.gz | GSE208337 | COVID-19 | |
| GSM6341276_B13_all_contig_annotations.csv.gz | GSE208337 | COVID-19 | |
| GSM6341277_B14_all_contig_annotations.csv.gz | GSE208337 | COVID-19 | |
| GSM6341278_B15_all_contig_annotations.csv.gz | GSE208337 | COVID-19 | |
| GSM6341279_B16_all_contig_annotations.csv.gz | GSE208337 | COVID-19 | |
| GSM6341280_B17_all_contig_annotations.csv.gz | GSE208337 | COVID-19 | |
| GSM6341281_B18_all_contig_annotations.csv.gz | GSE208337 | COVID-19 | |
| GSM6341282_B19_all_contig_annotations.csv.gz | GSE208337 | COVID-19 | |
| GSM6341283_B20_all_contig_annotations.csv.gz | GSE208337 | COVID-19 | |
| GSM6341284_B21_all_contig_annotations.csv.gz | GSE208337 | COVID-19 | |
| GSM6341285_B22_all_contig_annotations.csv.gz | GSE208337 | COVID-19 | |
| GSM6341286_B23_all_contig_annotations.csv.gz | GSE208337 | COVID-19 | |
| GSM6341287_B24_all_contig_annotations.csv.gz | GSE208337 | COVID-19 | |
| GSM6341288_B25_all_contig_annotations.csv.gz | GSE208337 | COVID-19 | |
| GSM6341289_B26_all_contig_annotations.csv.gz | GSE208337 | COVID-19 | |
| GSM6341290_B27_all_contig_annotations.csv.gz | GSE208337 | COVID-19 | |
| GSM6341291_B28_all_contig_annotations.csv.gz | GSE208337 | COVID-19 | |
| GSM6341292_B29_all_contig_annotations.csv.gz | GSE208337 | COVID-19 | |
| GSM6341293_B30_all_contig_annotations.csv.gz | GSE208337 | COVID-19 | |
| GSM6341294_B31_all_contig_annotations.csv.gz | GSE208337 | COVID-19 | |
| GSM6341295_B32_all_contig_annotations.csv.gz | GSE208337 | COVID-19 | |
| GSM6341296_B33_all_contig_annotations.csv.gz | GSE208337 | COVID-19 | |
| GSM6341297_B34_all_contig_annotations.csv.gz | GSE208337 | COVID-19 | |
| GSM6341298_B35_all_contig_annotations.csv.gz | GSE208337 | COVID-19 | |
| GSM6576371_S5_pbmc_BCR_before_filtered_contig_annotations.csv.gz | GSE213243 | Ovarian Cancer | |
| GSM6576374_S8_pbmc_BCR_after_filtered_contig_annotations.csv.gz | GSE213243 | Ovarian Cancer | 9 |
| hcd_1_filtered_contig_annotations.csv | GSE193867 | Healthy | |
| hcd_2_filtered_contig_annotations.csv | GSE193867 | Healthy | |
| hcd_3_filtered_contig_annotations.csv | GSE193867 | Healthy | |
| sle_1_filtered_contig_annotations.csv | GSE193867 | systemic lupus erythematosus | |
| sle_2_filtered_contig_annotations.csv | GSE193867 | systemic lupus erythematosus | |
| sle_3_filtered_contig_annotations.csv | GSE193867 | systemic lupus erythematosus | |
| filtered_contig_A_APP_annotations.csv | GSE193868 | Healthy | |
| filtered_contig_C_SPL_annotations.csv | GSE193868 | Healthy | 10 |

| | | | |
|---|---|---|---|
| filtered_contig_C_MLN_annotations.csv | GSE193868 | Healthy | |
| filtered_contig_C_APP_annotations.csv | GSE193868 | Healthy | |
| filtered_contig_B_SPL_annotations.csv | GSE193868 | Healthy | |
| filtered_contig_B_MLN_annotations.csv | GSE193868 | Healthy | |
| filtered_contig_B_APP_annotations.csv | GSE193868 | Healthy | |
| filtered_contig_A_SPL_annotations.csv | GSE193868 | Healthy | |
| filtered_contig_A_MLN_annotations.csv | GSE193868 | Healthy | |
| GSM5051382_AML1.BCR.all_contig_annotations.csv.gz | GSE165850 | AML | |
| GSM5051387_AML6.BCR.all_contig_annotations.csv.gz | GSE165850 | AML | |
| GSM5051386_AML5.BCR.all_contig_annotations.csv.gz | GSE165850 | AML | |
| GSM5051385_AML4.BCR.all_contig_annotations.csv.gz | GSE165850 | AML | |
| GSM5051384_AML3.BCR.all_contig_annotations.csv.gz | GSE165850 | AML | |
| GSM5051383_AML2.BCR.all_contig_annotations.csv.gz | GSE165850 | AML | 11 |
| GSM5231088_R125_filtered_contig_annotations.csv.gz | GSE171703 | COVID-19 | |
| GSM5231123_S65_filtered_contig_annotations.csv.gz | GSE171703 | COVID-19 | |
| GSM5231122_S50_filtered_contig_annotations.csv.gz | GSE171703 | COVID-19 | |
| GSM5231121_S407v2_filtered_contig_annotations.csv.gz | GSE171703 | COVID-19 | |
| GSM5231120_S407_filtered_contig_annotations.csv.gz | GSE171703 | COVID-19 | |
| GSM5231119_S356v2_filtered_contig_annotations.csv.gz | GSE171703 | COVID-19 | |
| GSM5231118_S356_filtered_contig_annotations.csv.gz | GSE171703 | COVID-19 | |
| GSM5231117_S33_filtered_contig_annotations.csv.gz | GSE171703 | COVID-19 | |
| GSM5231116_S281_filtered_contig_annotations.csv.gz | GSE171703 | COVID-19 | |
| GSM5231115_S272_filtered_contig_annotations.csv.gz | GSE171703 | COVID-19 | |
| GSM5231114_S266v2_filtered_contig_annotations.csv.gz | GSE171703 | COVID-19 | |
| GSM5231113_S266_filtered_contig_annotations.csv.gz | GSE171703 | COVID-19 | |
| GSM5231112_S218_filtered_contig_annotations.csv.gz | GSE171703 | COVID-19 | |
| GSM5231111_S210v2_filtered_contig_annotations.csv.gz | GSE171703 | COVID-19 | |
| GSM5231110_S201_filtered_contig_annotations.csv.gz | GSE171703 | COVID-19 | |
| GSM5231109_S130_filtered_contig_annotations.csv.gz | GSE171703 | COVID-19 | |
| GSM5231108_S92_filtered_contig_annotations.csv.gz | GSE171703 | COVID-19 | |
| GSM5231107_S609_filtered_contig_annotations.csv.gz | GSE171703 | COVID-19 | |
| GSM5231106_S586_filtered_contig_annotations.csv.gz | GSE171703 | COVID-19 | |
| GSM5231105_S564_filtered_contig_annotations.csv.gz | GSE171703 | COVID-19 | |
| GSM5231104_S537_filtered_contig_annotations.csv.gz | GSE171703 | COVID-19 | |
| GSM5231103_S48_filtered_contig_annotations.csv.gz | GSE171703 | COVID-19 | |
| GSM5231102_S376_filtered_contig_annotations.csv.gz | GSE171703 | COVID-19 | |
| GSM5231101_S305_filtered_contig_annotations.csv.gz | GSE171703 | COVID-19 | |
| GSM5231100_S24_filtered_contig_annotations.csv.gz | GSE171703 | COVID-19 | |
| GSM5231099_S214_filtered_contig_annotations.csv.gz | GSE171703 | COVID-19 | 12 |

| | | | |
|---|---|---|---|
| GSM5231098_S210_filtered_contig_annotations.csv.gz | GSE171703 | COVID-19 | |
| GSM5231097_S20_filtered_contig_annotations.csv.gz | GSE171703 | COVID-19 | |
| GSM5231096_S171_filtered_contig_annotations.csv.gz | GSE171703 | COVID-19 | |
| GSM5231095_S166_filtered_contig_annotations.csv.gz | GSE171703 | COVID-19 | |
| GSM5231094_S155_filtered_contig_annotations.csv.gz | GSE171703 | COVID-19 | |
| GSM5231093_S144_filtered_contig_annotations.csv.gz | GSE171703 | COVID-19 | |
| GSM5231092_S116_filtered_contig_annotations.csv.gz | GSE171703 | COVID-19 | |
| GSM5231091_R6_filtered_contig_annotations.csv.gz | GSE171703 | COVID-19 | |
| GSM5231090_R478910_filtered_contig_annotations.csv.gz | GSE171703 | COVID-19 | |
| GSM5231089_R3_filtered_contig_annotations.csv.gz | GSE171703 | COVID-19 | |
| GSM4633683_T11_BCR_filtered_contig_annotations.csv.gz | GSE150825 | Nasopharyngeal tumor | |
| GSM4633691_T9_BCR_filtered_contig_annotations.csv.gz | GSE150825 | Nasopharyngeal lymphoid hyperplasia | |
| GSM4633690_T7_BCR_filtered_contig_annotations.csv.gz | GSE150825 | Nasopharyngeal lymphoid hyperplasia | |
| GSM4633689_T5_BCR_filtered_contig_annotations.csv.gz | GSE150825 | Nasopharyngeal lymphoid hyperplasia | |
| GSM4633688_T215_BCR_filtered_contig_annotations.csv.gz | GSE150825 | Nasopharyngeal tumor | |
| GSM4633687_T8_BCR_filtered_contig_annotations.csv.gz | GSE150825 | Nasopharyngeal tumor | |
| GSM4633686_T6_BCR_filtered_contig_annotations.csv.gz | GSE150825 | Nasopharyngeal tumor | |
| GSM4633685_T147_BCR_filtered_contig_annotations.csv.gz | GSE150825 | Nasopharyngeal tumor | |
| GSM4633684_T4_BCR_filtered_contig_annotations.csv.gz | GSE150825 | Nasopharyngeal tumor | 13 |
| GSM3576459_C33_pBMC_full-length_productive_BCR_table.tsv.gz | GSE125527 | Healthy | |
| GSM3576432_C9_R_full-length_productive_BCR_table.tsv.gz | GSE125527 | Healthy | |
| GSM3576433_C12_R_full-length_productive_BCR_table.tsv.gz | GSE125527 | Healthy | |
| GSM3576434_C16_R_full-length_productive_BCR_table.tsv.gz | GSE125527 | Healthy | |
| GSM3576435_U4_R_full-length_productive_BCR_table.tsv.gz | GSE125527 | Ulcerative Colitis | |
| GSM3576436_U5_R_full-length_productive_BCR_table.tsv.gz | GSE125527 | Ulcerative Colitis | |
| GSM3576437_U34_R_full-length_productive_BCR_table.tsv.gz | GSE125527 | Ulcerative Colitis | |
| GSM3576438_U35_R_full-length_productive_BCR_table.tsv.gz | GSE125527 | Ulcerative Colitis | |
| GSM3576439_U41_R_full-length_productive_BCR_table.tsv.gz | GSE125527 | Ulcerative Colitis | |
| GSM3576440_U45_R_full-length_productive_BCR_table.tsv.gz | GSE125527 | Ulcerative Colitis | |
| GSM3576441_C17_R_full-length_productive_BCR_table.tsv.gz | GSE125527 | Healthy | |
| GSM3576442_C18_R_full-length_productive_BCR_table.tsv.gz | GSE125527 | Healthy | |
| GSM3576443_C19_R_full-length_productive_BCR_table.tsv.gz | GSE125527 | Healthy | |
| GSM3576444_C21_R_full-length_productive_BCR_table.tsv.gz | GSE125527 | Healthy | 14 |

| | | | |
|---|---|---|---|
| GSM3576445_C30_R_full-length_productive_BCR_table.tsv.gz | GSE125527 | Healthy | |
| GSM3576446_C12_pBMC_full-length_productive_BCR_table.tsv.gz | GSE125527 | Healthy | |
| GSM3576447_C16_pBMC_full-length_productive_BCR_table.tsv.gz | GSE125527 | Healthy | |
| GSM3576448_U4_pBMC_full-length_productive_BCR_table.tsv.gz | GSE125527 | Ulcerative Colitis | |
| GSM3576449_U5_pBMC_full-length_productive_BCR_table.tsv.gz | GSE125527 | Ulcerative Colitis | |
| GSM3576450_U34_pBMC_full-length_productive_BCR_table.tsv.gz | GSE125527 | Ulcerative Colitis | |
| GSM3576451_U35_pBMC_full-length_productive_BCR_table.tsv.gz | GSE125527 | Ulcerative Colitis | |
| GSM3576452_U41_pBMC_full-length_productive_BCR_table.tsv.gz | GSE125527 | Ulcerative Colitis | |
| GSM3576453_U44_pBMC_full-length_productive_BCR_table.tsv.gz | GSE125527 | Ulcerative Colitis | |
| GSM3576454_U45_pBMC_full-length_productive_BCR_table.tsv.gz | GSE125527 | Ulcerative Colitis | |
| GSM3576455_C17_pBMC_full-length_productive_BCR_table.tsv.gz | GSE125527 | Healthy | |
| GSM3576456_C18_pBMC_full-length_productive_BCR_table.tsv.gz | GSE125527 | Healthy | |
| GSM3576457_C21_pBMC_full-length_productive_BCR_table.tsv.gz | GSE125527 | Healthy | |
| GSM3576458_C30_pBMC_full-length_productive_BCR_table.tsv.gz | GSE125527 | Healthy | |
| trust4_tcga_deidentified_report.tsv | NA | Cancer | [15] |
| GSM4785642_P6_BAL_BCR.filtered_contig_annotations.csv.gz | GSE158038 | COVID-19 | |
| GSM4785643_P17_Bal1_BCR.filtered_contig_annotations.csv.gz | GSE158038 | COVID-19 | |
| GSM4785644_P17_Bal2_BCR.filtered_contig_annotations.csv.gz | GSE158038 | COVID-19 | [16] |
| GSE215802_BCR_filtered_contig_annotations_06.csv.gz | GSE215802 | COVID-19 | |
| GSE215802_BCR_filtered_contig_annotations_08.csv.gz | GSE215802 | COVID-19 | |
| GSE215802_BCR_filtered_contig_annotations_07.csv.gz | GSE215802 | COVID-19 | NA |
| GSM6176194_filtered_contig_annotations_p1_biopsy1_bcr.csv.gz | GSE203552 | CNS Lymphoma | |
| GSM6176211_filtered_contig_annotations_p2_blood_bcr.csv.gz | GSE203552 | CNS Lymphoma | |
| GSM6176209_filtered_contig_annotations_p2_biopsy_bcr.csv.gz | GSE203552 | CNS Lymphoma | |
| GSM6176206_filtered_contig_annotations_p1_csf_bcr.csv.gz | GSE203552 | CNS Lymphoma | |
| GSM6176204_filtered_contig_annotations_p1_blood3_bcr.csv.gz | GSE203552 | CNS Lymphoma | |
| GSM6176202_filtered_contig_annotations_p1_blood2_bcr.csv.gz | GSE203552 | CNS Lymphoma | |
| GSM6176200_filtered_contig_annotations_p1_blood1_bcr.csv.gz | GSE203552 | CNS Lymphoma | |
| GSM6176198_filtered_contig_annotations_p1_biopsy3_bcr.csv.gz | GSE203552 | CNS Lymphoma | |
| GSM6176196_filtered_contig_annotations_p1_biopsy2_bcr.csv.gz | GSE203552 | CNS Lymphoma | [17] |
| GSE203610_FL_meta_bcr.csv.gz | GSE203610 | Follicular Lymphoma | [18] |
